# Reconstructing parent genomes using siblings and other relatives

**DOI:** 10.1101/2024.05.10.593578

**Authors:** Ying Qiao, Ethan M. Jewett, Kimberly F. McManus, William A. Freyman, Joanne E. Curran, Sarah Williams-Blangero, John Blangero, The 23andMe Research Team, Amy L. Williams

## Abstract

Reconstructing the DNA of ancestors from their descendants has the potential to empower phenotypic analyses (including association and genetic nurture studies), improve pedigree reconstruction, and shed light on the ancestral population and phenotypes of ancestors. We developed HAPI-RECAP, a method that reconstructs the DNA of parents from full siblings and their relatives. This tool leverages HAPI2’s output, a new phasing approach that applies to siblings (and optionally one or both parents) and reliably infers parent haplotypes but does not link the ungenotyped parents’ DNA across chromosomes or between segments flanking ambiguities. By combining IBD between the reconstructed parents and the relatives, HAPI-RECAP resolves the source parent of these segments. Moreover, the method exploits crossovers the children inherited and sex-specific genetic maps to infer the reconstructed parents’ sexes. We validated these methods on research participants from both 23andMe, Inc. and the San Antonio Mexican American Family Studies. Given data for one parent, HAPI2 reconstructs large fractions of the missing parent’s DNA, between 77.6% and 99.97% among all families, and 90.3% on average in three- and four-child families. When reconstructing both parents, HAPI-RECAP inferred between 33.2% and 96.6% of the parents’ genotypes, averaging 70.6% in four-child families. Reconstructed genotypes have average error rates < 10^−3^, or comparable to those from direct genotyping. HAPI-RECAP inferred the parent sexes 100% correctly given IBD-linked segments and can also reconstruct parents without any IBD. As datasets grow in size, more families will be implicitly collected; HAPI-RECAP holds promise to enable high quality parent genotype reconstruction.

## Introduction

Parents transmit half their DNA to each child, with recombination and independent segregation leading to distinct genetic inheritance among multiple offspring and the potential to reconstruct a parent’s genome from these offspring’s DNA. Applications of parent DNA reconstruction are vast, including increasing power in linkage or genome-wide association studies (GWAS) by adding data from ungenotyped individuals ^1,2^. Such augmented data expands on the benefits of GWAS by proxy (i.e., analyzing a child in lieu of a proband) ^3,4^ in that it isolates each parent’s DNA and, given multiple children, should typically include more than 50% of it. Moreover, parents’ DNA is essential to studies of indirect genetic effects—so-called genetic nurture, or the association of parents’ untransmitted alleles with a child’s phenotype ^5^—which has motivated the development of parent DNA reconstruction methods ^6,7^. Other applications of reconstructed ancestor DNA include improving the accuracy of relatedness inference ^8^ and therefore also of pedigree reconstruction ^9–11^, and revealing the population of origin of those ancestors ^12^ together with some of their traits.

Given a pedigree that lacks inbreeding, one approach to reconstructing ancestors’ DNA involves simply placing identity-by-descent (IBD) segments—i.e., long stretches of identical DNA that two or more individuals inherited from a common ancestor—on the ancestors that connect those IBD carriers ^1^. For example, the IBD segments two first cousins share are necessarily also carried by their two parents (who are full siblings of one another). This type of reconstruction becomes more complex in small or endogamous populations and in inbred pedigrees. In these cases, the ancestor that transmitted the IBD segment may be difficult to decipher, but recent work has made progress in these settings ^13^. Even when analyzing individuals from large, outbred populations, placing IBD segments on an ancestral couple using their descendants’ DNA—e.g., the grandparents of two first cousins—requires IBD sharing to other relatives. This is necessary to disambiguate which ancestor each segment descends from and typically requires non-descendant relatives of the couple, e.g., second cousins of the first cousins. Note that this ambiguity also applies to the reconstruction of both parents using full siblings.

Depending on the pedigree structure and available data, not all of an ancestor’s transmitted DNA will be IBD between two or more genotyped descendants, which limits the amount of the ancestor’s genome the IBD-placing approach can reconstruct. In specific settings, if an ancestor’s DNA can be detected by other means, it is possible to avoid this reliance on IBD segments. One striking example is Hans Jonatan, the child of an African mother and European father who lived in Iceland at a time when few others had African ancestry. Jonatan’s descendants harbor segments of his maternal genome and use of an extensive genealogy together with local ancestry inference enabled the reconstruction of 38% of his maternal DNA (or ∼19% of Jonatan’s genome) from 182 descendants ^12^.

This paper focuses on reconstructing the DNA of parents from a set of siblings, which affords the opportunity to reconstruct considerably larger fractions of their genomes even with fairly small amounts of IBD sharing to other relatives. This is because of the highly constrained sources of genetic material in full siblings— two diploid parents subject to recombination only between their own pair of haplotypes. While each child inherits only half of the genomes of each parent, randomized transmissions lead *c* children to inherit an average proportion of 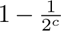 of both parents’ genomes. Notably, while most approaches to reconstruct DNA from deeper ancestors require a pedigree structure, which may contain errors, identifying full siblings and parent-child pairs is feasible with >99% recall and few false positives ^14^, enabling highly accurate nuclear family detection.

Existing methods to reconstruct parent genotypes from sibling data include those designed to study indirect genetic effects ^6,7^ and many others that implement family-based phasing algorithms ^15–17^. The latter are the most similar to our approach, but a fundamental challenge in this problem is determining which parent each genotype belongs to. Methods designed for studying indirect genetic effects do not attempt to distinguish the two parents: these analyses do not require linking the parents’ genotypes across loci or even knowing which parent (father or mother) carries each genotype ^6,7^. By tracking which haplotype each parent transmitted at each marker (i.e., the inheritance vector ^17^), family-based phasers can effectively link parent DNA at neighboring sites. Yet without auxiliary information, they cannot assign DNA from distinct chromosomes to the same parent and are subject to ambiguities that can conflate the two parents’ DNA on the same chromosome. Indeed, the scoring metric for one recent method allowed the reconstructed parent genotypes to correspond to either parent arbitrarily at each marker ^15^. Analyzing a pedigree that includes data for non-descendant relatives of the parents can help separate the parents’ DNA, but many pedigree phasing methods use computationally expensive Markov chain Monte Carlo (MCMC) approaches that require marker thinning to those in linkage equilibrium and (depending on the parameters used) may yield lower accuracy genotypes than we seek ^16^.

Here, we introduce an approach to reconstruct the genotypes of parents from full siblings using a combination of family-based phasing and signals from both IBD sharing and sex-specific genetic maps ^18,19^ to disambiguate which DNA belongs to each parent. First, HAPI2, an updated and expanded version of HAPI ^20^, infers minimum-recombinant haplotypes in nuclear families, applying to a set of full siblings alone or to siblings and one or both of their parents. When genotype data for one or both parents is missing, HAPI2 enables joint phasing of the siblings and infers haplotype segments of the missing parents using only the children’s genotype data. This joint phasing has low error and, depending on the number of children, provides long haplotypes for the parents, even up to chromosome-scale. Still, as mentioned above, these reconstructed data have ambiguous linkage to the two parents when segments occur on different chromosomes and, in cases of ambiguous regions, even distinct segments on the same chromosome (see Methods).

To assign the reconstructed DNA to the parents, our new tool HAPI-RECAP (REConstruct Ancestral genotyPes) leverages IBD between HAPI2’s reconstructed parent haplotypes and relatives of the children. The basis of this inference is that, if a relative is connected to the children only through one parent, that relative’s IBD segments must be shared with the DNA that belongs to that parent. Using only one IBD segment, HAPI-RECAP infers not only the genotypes of the related parent, but further deduces that the homologous reconstructed DNA belongs to the other parent (see Figure 1B). Importantly, HAPI2’s reconstructed DNA may span a much larger interval than the IBD segment, but that one segment suffices to link the full reconstructed interval to the parents. This analysis is similar to performing inter-chromosomal phasing using IBD sharing to relatives ^21–23^: even though HAPI-RECAP analyzes IBD to reconstructed parent DNA, the goal of determining the source parent of the DNA and the reliance on individuals related to the siblings through only one lineage equally applies to both problems.

**Figure 1:**
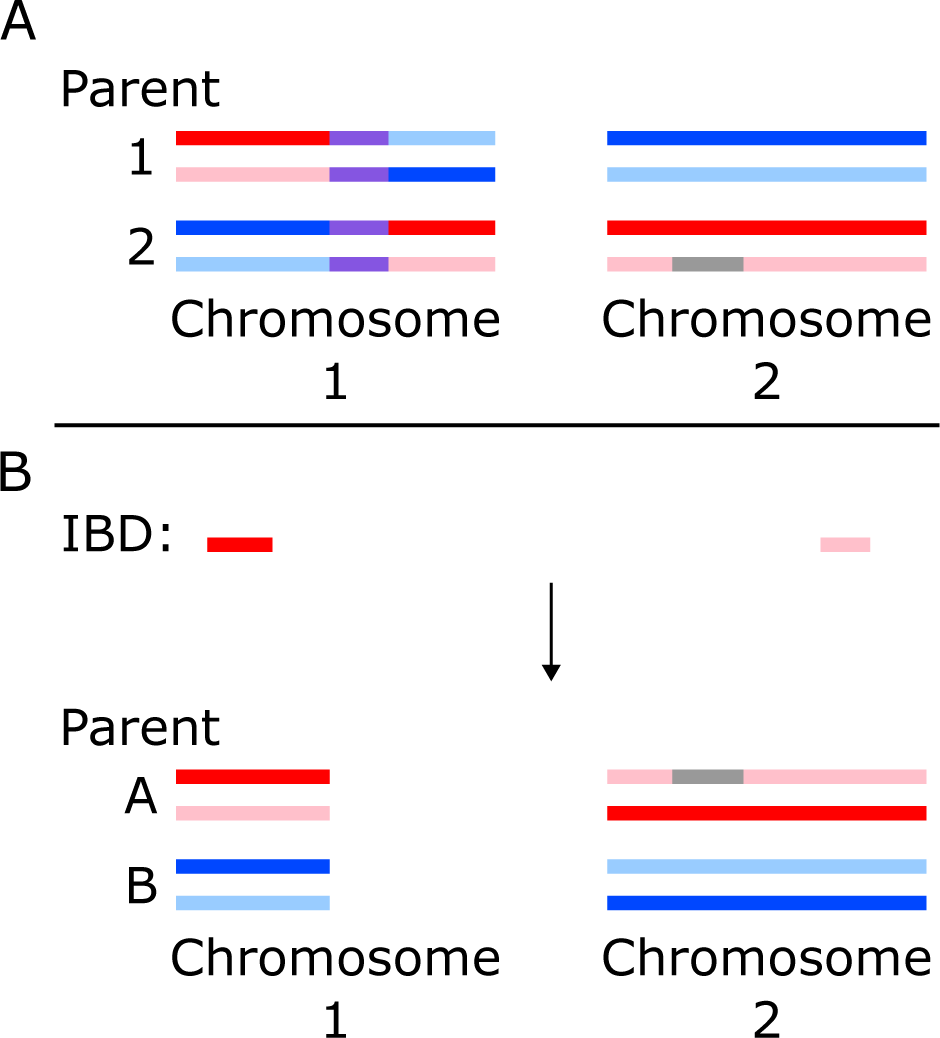
When analyzing only siblings, HAPI2 produces reconstructed parent genomes with mixed maternal and paternal origin and HAPI-RECAP infers linkage between reconstructed segments using IBD to relatives. The example depicts two chromosomes with maternal haplotypes shown in red and pink and paternal haplotypes in dark and light blue. Regions in which the two parents’ haplotype transmissions are indistinguishable are shown in purple. These intervals prevent linkage of the DNA on either side of them to the same parent, so the flanking maternal and paternal DNA may be assigned to different genomes. Gray intervals represent a non-reconstructed haplotype, which occur when a parent transmits only one haplotype to the children; linkage is retained across these intervals. (A) Example output from HAPI2, which infers reconstructed segments that separate the parents’ DNA, shown here with contiguous non-purple color horizontally. The parents’ haplotype transmissions are ambiguous near the center of chromosome 1 (purple), and the reconstructed DNA in parent 1’s genome is maternal to the left of this interval and paternal to the right. Parent assignments to the genomes labeled parent 1 and parent 2 are arbitrary across chromosomes. (B) Two IBD segments shared between the reconstructed parent DNA and a maternal relative are shown with horizontal position aligned to the reconstructed genomes in (A). These segments allow assignment of the left reconstructed segment on chromosome 1 and the reconstructed segment on chromosome 2 (which spans the entire chromosome) to the two parents. As the right reconstructed segment on chromosome 1 has no IBD segment covering it, it is not assigned to the parents. This paper labels the unsexed reconstructed parents HAPI-RECAP produces as parents A and B.

We further incorporated signals from genetic maps into HAPI-RECAP by calculating the probability that a male or female parent transmitted the crossovers HAPI2 detects in the children. Recent work showed that the profound differences between male and female genetic maps are sufficient to detect the sex of the shared parent of two half-siblings (or the parent linking a grandparent-grandchild pair) using autosomal IBD alone ^24^. Whereas all IBD segments (genome-wide) between half-siblings can be attributed to one parent, HAPI2’s reconstructed segments are initially unlinked. As such, the genetic map-based inference is most powerful after using IBD to assign several reconstructed segments to the two (unsexed) parents: then the method can jointly consider the crossovers in each segment to infer the parents’ sexes. Even so, we show that some individual reconstructed segments are sufficiently informative to also allow this inference. Overall, this component of HAPI-RECAP enables inference of parent sexes even if no relative shares IBD to the X chromosome; it also provides parent-of-origin phasing of the children even when both parents are ungenotyped.

Finally, in addition to HAPI-RECAP’s reconstruction of both parents using DNA from siblings and their relatives, HAPI2 alone reconstructs the DNA of one parent given data for siblings and their other parent (and no other relatives). The abundant information for this problem makes the reconstruction highly complete, with diploid reconstruction across much of the DNA and haploid reconstruction at loci in which the parent transmitted only one haplotype to the children (which happens at a rate of 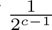 given *c* children).

We tested these methods on nuclear families composed of research participants from 23andMe, Inc. and the San Antonio Mexican American Family Studies (SAMAFS) ^25–27^. To allow comparison of the reconstructed parents with direct genotypes, we restricted to families with data available for both parents. The validation includes two scenarios: reconstructing one parent using three or more children and data from the other parent, and reconstructing both parents using four or more children and their relatives. We also present results from using the crossovers HAPI2 detects in the siblings to infer the sexes of the parents and for using these crossovers and no relatives to reconstruct both parents.

## Methods

Our approach to reconstructing parents’ DNA has three key components. First, HAPI2 jointly phases a set of siblings (or siblings and one of their parents) using a hidden Markov model (HMM) that reconstructs large fractions of the parents’ DNA and builds a one-generation ancestral recombination graph (ARG). While effective, in cases where both parents’ data are missing, this does not infer which reconstructed segments on different chromosomes belong to the same parent; linkage is also impossible for HAPI2 to track across ambiguous regions on the same chromosome (Figure 1A). To resolve these ambiguities, HAPI-RECAP, the second component, uses IBD segments the parents share with other relatives to link reconstructed DNA to the same parent (Figure 1B). Finally, HAPI-RECAP also leverages the crossovers that HAPI2 detects to infer the sex of the parent that transmitted one or more of the reconstructed segments. This last component enables assignment of some reconstructed segments to the parents even with no overlapping IBD segment, enables sex assignment of the reconstructed parents, and thereby determines parent-of-origin of some of the DNA the siblings inherited.

We describe each of these components and our data processing below. Throughout this paper, we refer to *reconstructed segments* as contiguous stretches of reconstructed parent DNA that HAPI2 outputs when the transmission patterns allow the two parents’ DNA to be distinguished. (These are segments of uninterrupted red, pink, and dark or light blue color that may include a gray region in Figure 1A.)

### Jointly phasing siblings

We developed HAPI2, which extends the nuclear family phasing method HAPI ^20^ to cases where one or both parents’ data are missing, adds an error model, and has a new codebase. Both HAPI and HAPI2 implement an HMM, though HAPI2 uses recombination counts and not probabilities for defining the likelihood of different phases, and it chooses the Viterbi decoding to obtain minimum-recombinant phase. (HAPI implements both minimum-recombinant and maximum-likelihood phase, though in practice these are often similar ^20^).

HAPI generates states based on the genotypes of the parents and HAPI2 uses a similar logic but incorporates states corresponding to all possible parent genotypes, as dictated by the children’s genotypes. For example, at biallelic markers, if one child is homozygous for allele A and another for allele B, then, assuming no genotyping error, the parents must both be heterozygous (A/B); alternatively, if the children have only one homozygous genotype (say for allele A) and heterozygous calls, this could correspond to one parent being heterozygous and the other homozygous for A or could arise when both parents are heterozygous. In biallelic data, most markers provide incomplete information about the parents’ genotypes and the children’s phase. Yet HAPI2’s states include an inheritance vector ^17^ of length 2*c*, where *c* is the number of children, which specifies which haplotype each parent transmitted to each child at the corresponding marker. Recombinations are encoded as changes in the inheritance vector between sites.

HAPI2’s algorithm proceeds as follows, and we provide details in the subsections below. The method analyzes markers one by one in physical order, and, at each marker, HAPI2 first detects the set of all possible parent genotypes consistent with the observed children’s genotypes (incorporating any available parent data, including both parents’ genotypes if available; see A.1: Detecting marker type in the Appendix). Next, HAPI2 generates partial states (i.e., deriving only from the data at the current marker) for each of the possible parent genotypes at the marker (see A.2: Generating partial states). The method then builds full states by mapping all states at the previous marker to each partial state, filling in any undefined inheritance vector values in the partial state (see A.4: Generating full states). When both parents’ data are missing, in some cases the children’s genotypes alone cannot distinguish which parent transmitted each haplotype, even with the neighboring inheritance vectors, and HAPI2 flags these parent ambiguities (see A.5: Distinguishing parent genotypes). Additionally, if one or both parents are missing data, HAPI2 infers otherwise latent recombinations that result in all children inheriting only one of the parent’s haplotypes (this is necessary to avoid incorrectly assuming the parent is homozygous in such a region; see A.6: Inferring that a parent transmitted only one haplotype). HAPI2 supports X chromosome phasing, which involves some specific considerations (see A.7: X chromosome phasing). The method includes error detection (see A.8: Detecting genotyping errors) and several optimizations that make it efficient to apply in practice even though the size of the state space is *O*(2^2c^) and a naive implementation would be prohibitive for analyzing large families (see A.9: Optimizations). After analyzing all markers on a chromosome, HAPI2’s last step is back tracing to resolve the phase and parent haplotypes according to the globally minimum-recombinant state path (see A.10: Back tracing).

### Using IBD to resolve the parent assignment of reconstructed segments

As described above, when analyzing only siblings, HAPI2 outputs a number of reconstructed parent segments, and which parent these belong to is *a priori* unclear (Figure 1A). HAPI-RECAP uses IBD between these reconstructed segments and the siblings’ relatives to resolve parental origin (Figure 1B). The approach assumes that the relationship to the siblings is only through one parent, which is likely for many close relatives in most populations. Notably, however, individuals that descend from both parents and some forms of inbreeding violate this assumption; HAPI-RECAP takes steps to detect such individuals (below) and does not use them in its analysis.

HAPI-RECAP works by considering all IBD segments each relative shares with either parent 1 or 2. Recall that although HAPI2 outputs the DNA using two parents, the true parental origin is not known, and, in fact, the parents’ DNA is mixed between these individuals (Figure 1A). Given the reconstructed phased parent data from HAPI2 and phased data for the siblings’ relatives, a preliminary step before running HAPI-RECAP is to detect IBD segments using TPBWT ^28^ (or another IBD detector), and we did this with phase correction disabled.

HAPI-RECAP takes these IBD segments as input and then performs the following steps:

1. Merge overlapping IBD segments any relative *r* shares with a parent *p* on both haplotypes. (If the parent had only one haplotype reconstructed, HAPI2’s output VCF represents this with diploid homozygous calls—half-missing calls violate the VCF specifications—and we are only interested in knowing that there is ≥ 1 IBD segment to the parent at the locus.)
2. Delete overlapping IBD at locations a relative *r* shares with both parents (retaining any part of the segments that do not overlap). In regions where the two parents’ haplotype transmissions are indistinguishable (see A.5: Distinguishing parent genotypes), the output VCF includes only uninformative sites and those where both parents are heterozygous, so the reconstructed DNA for the two parents do not differ.
3. Retain only > 9 cM long IBD segments.
4. Sum the length of remaining IBD and retain only close relatives, i.e., those sharing > 600 cM combined to the two parents.
5. Filter any relative *r* with sufficient evidence of sharing IBD with both parents, i.e., where *n_r,b_/n_r_ >* 0.1, with *n_r_* being the number of reconstructed parent segments *r* shares ≥ 1 IBD segment with (to either parent), and *n_r,b_ ≤ n_r_* is the number of reconstructed segments *r* has IBD sharing to both parents in. In this step and below, an IBD segment must overlap a reconstructed segment by > 6 cM to be considered.
6. Discard reconstructed segments in which any remaining relative *r* shares IBD to both parents (counted in *n_r,b_*above). Depending on the inheritance vectors and recombinations, HAPI2 occasionally inadvertently swaps the parent labels within a reconstructed segment. This can occur when the two parents’ inheritance vector elements are similar and a recombination transmitted by one parent can be modeled with zero recombinations by swapping the parent labels. (Notably HAPI2 lacks the data to detect that this is happening and makes this assignment because it yields minimum-recombinant phase.)
7. Sum *L_r_*, the cM length of the reconstructed segments each relative *r* shares an IBD segment to.
8. For every pair of relatives *r*_1_*, r*_2_, count the number of reconstructed segments in which both share an IBD segment to the same parent *s_r_*_1_ *_,r_*_2_ and opposite parents *o_r_*_1_ *_,r_*_2_.
9. Let 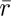 = arg max*_r_ L_r_* and, considering only relatives *r* where 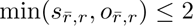, (i.e., 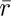 and *r* share IBD segments consistently to either the same or opposite parents in nearly all overlapping reconstructed segments), calculate a score 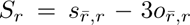 if 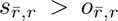 and 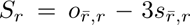 otherwise. Let *r′* = arg max*_r_ S_r_*, the relative with the highest score for sharing IBD to either the same or opposite parent as 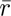.
10. Reconstruct the two parents A and B using 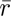 and *r′* by assigning reconstructed segments 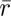 shares IBD in to parent A and the homologous reconstructed segment HAPI2 assigned to the opposite parent to parent B. Assign reconstructed segments *r′* shares IBD in to parent A if 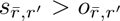 and to parent B otherwise; assign homologous segments to the opposite parent. Discard any reconstructed segments in which both 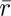 and *r′* share IBD segments but with conflicting signals about which parent the segments should be assigned to.

### Using sex-specific maps to infer parent sex

HAPI2 detects crossovers the parents transmitted to a set of children, and, when neither parents’ data are available, HAPI-RECAP uses the locations of these crossovers to infer the sexes of the reconstructed parents. Additionally, if no IBD covers a reconstructed segment, HAPI-RECAP can still assign it to a parent if the crossovers are sufficiently informative.

The inference proceeds by analyzing all crossovers in a reconstructed segment or in all the reconstructed segments assigned to parents A and B using IBD. In both cases there are two sets of crossovers, each set transmitted by the same parent, and we will refer to the transmitting parents as parent 1 and 2, which have unknown sex. The model assumes the crossovers are independent (as in CREST ^24^) and leverages the genetic lengths of each crossover interval (Figure 2). That is, crossovers are detectable in an interval bounded on either side by a heterozygous SNP; because not all sites are heterozygous, the basepair position of the crossover is not observable (moreover, in this paper we analyze SNP array genotypes).

**Figure 2:**
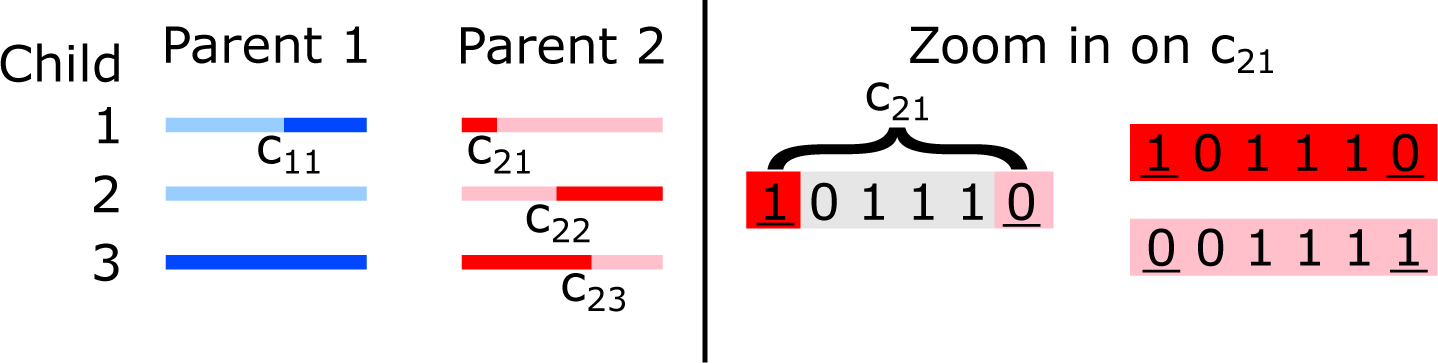
HAPI-RECAP uses crossover intervals from two unsexed parents and sex-specific genetic maps to infer the parents’ sexes. The example depicts colored haplotype segments parents 1 and 2 transmitted to three children, with switches in color along the horizontal haplotypes corresponding to crossovers. Parent 1 has a set of crossover intervals *C*_1_, which for this segment contains only one interval, *c*_11_; parent 2 has three crossover intervals in *C*_2_. In general, crossovers can only be localized to an interval bounded on each side by a heterozygous site. The right panel shows a zoomed in example for one crossover interval. Here, each marker has alleles 0 and 1 and we show the parental haplotypes in red and pink at six consecutive SNPs, with heterozygous SNPs underlined. The interval *c*_21_ spans these six SNPs, though it is an open interval on both sides and excludes the heterozygous sites. The transmitted haplotype (red or pink) cannot be determined inside the interval because the SNPs are homozygous.

Let *ℓ_s_*(*c*) be the genetic length (here in Morgans) of a given crossover interval *c* in the genetic map ^19^ for sex *s ∈ {*F, M} (F for female, M for male). If this map length is 0, HAPI-RECAP instead assigns *ℓ_s_*(*c*) = 10^−6^. Then using the Poisson probability *Pr* (*k*; *ℓ*) of *k* events occurring in an interval of length *ℓ*, let *C*_1_ and *C*_2_ be the sets of crossovers transmitted by parents 1 and 2, respectively, and

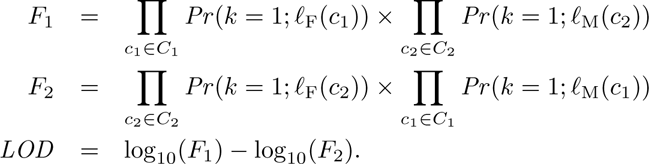

Then *LOD* is the log_10_-odds of parent 1 versus parent 2 being female: *LOD >* 0 suggests that parent 1 is female and *LOD <* 0 suggests parent 2 is.

To be conservative, HAPI-RECAP only assigns a reconstructed segment to a parent if |*LOD* | > 3. Typically, when considering many reconstructed segments jointly (i.e., after using IBD to reconstruct parent A and B), |*LOD | ≫* 3 (Results). The interval length, i.e., distance between flanking informative sites, can be large for crossovers at the beginning of a region with one haplotype transmitted (A.6: Inferring that a parent transmitted only one haplotype) and near the boundaries of regions in which the two parents’ haplotype transmissions are indistinguishable (A.5: Distinguishing parent genotypes). To avoid noise from such poorly localized crossovers, HAPI-RECAP requires that the crossover intervals it analyzes are contained inside the reconstructed segment and are fewer than 45 markers long.

### Validation and quality metrics for reconstructed genotypes

To evaluate HAPI2’s reconstruction of one parent and HAPI-RECAP’s reconstruction of both parents, we compared the reconstructed genotypes with the available genotype data for each parent and calculated two primary metrics: the amount of DNA reconstructed and the error rate. We calculate the amount of DNA reconstructed as 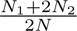, where *N*_1_ and *N*_2_ are the number of sites reconstructed with one copy and both copies, respectively. *N* is the total number of sites in the directly genotyped parent data that HAPI2 phased when we ran it on the complete nuclear family (i.e., it excludes sites HAPI2 called ambiguous or as having Mendelian or other errors). We call the latter data the truth set. The error rate quantifies the rate of incorrect reconstructed sites and we define this as 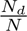, where *N_d_* the number of sites with genotypes that differ from the the truth set. We count any difference in genotype at sites HAPI2 reconstructs on both copies as an error, while sites that are reconstructed on one copy need only match one allele in the truth set. Thus, while we calculate the amount of DNA reconstructed using both parental haplotypes, even one allelic difference at a site reconstructed on both haplotypes counts as an error.

### Data processing

We analyzed SNP array data from 114 nuclear families contained in the SAMAFS pedigrees. These analyses started from a dataset that had extensive quality control filters carried out in a previous study ^29^, including: mapping the SNP array probe sequences to GRCh37; removing SNPs with ≥1% missingness and those that failed a test for differences between male and female allele frequencies; and removing samples with greater than 2% missing sites. Since monozygotic (MZ) twins share the same genotypes, we also removed six individuals to keep only one member of each MZ twin pair. The final dataset includes 2,456 individuals and 511,007 SNPs. Among these individuals, there are 114 nuclear families that include genotype data for both parents and between 3 to 12 children (average 4.5 children).

The 23andMe research participants include thousands of nuclear families, and we identified those with data for both parents and four or more children and then randomly sampled to obtain 220 families in total (100 4-child, 60 5-child, 40 6-child, 15 7-child, two 8-child, and three 10-child families). To ascertain these families, we considered research participants genotyped on the most recent 23andMe SNP array (the ‘v5’ array). We identified potential MZ twins or sample duplicates as pairs that share IBD ^30^ on both chromosomes (IBD2) totaling more than 3,400 cM, creating sets in which all individuals share IBD at this level to another member of the set, and retained only one arbitrarily chosen individual from each set. We then detected putative parent-child pairs as those sharing >3,400 cM of IBD but <3,400 cM of IBD2 and considered the individual with the older self-reported date of birth as the putative parent. After filtering out samples with >1 putative mother, we ascertained mothers and fathers that have an overlapping set of ≥4 children; these parents together with their children constitute the families from which we sampled study subjects.

We ran HAPI2 on each of these nuclear families while censoring the parent data (either both or one parent, depending on the analysis) and with error detection options of --err_min 1 and --err_dist 5. HAPI2 detects crossovers by scanning the inferred inheritance vectors sequentially: an informative marker *m* where an inheritance vector element for some child changes relative to the previous site indicates a potential crossover, and the interval it occurred in is from the most recent upstream informative site to *m*. To filter errors or non-crossover gene conversions, the method requires a user-specified number of downstream informative sites to retain the new inheritance vector value. For our analyses, we considered a change in inheritance vector that spans at least 10 informative sites to be a crossover and so ran with --detect_co 10.

To detect IBD segments in the reconstructed SAMAFS parents, we ran Beagle 5.4 (22jul22.46e)^31^ on the SAMAFS data with parents removed and then ran TPBWT ^28^ with phase correction disabled on the reconstructed parent haplotypes output by HAPI2 and these Beagle-phased individuals.

The relatives of each 23andMe family that we used for detecting IBD with the reconstructed parents are research participants who share ≥600 cM and ≤3,000 cM of IBD and ≤20 cM of IBD2 with at least one child in any family. As part of a separate study, 23andMe researchers used SHAPEIT 5.0.0^32^ to phase more than 8.6 million research-consented individuals ^23^and we extracted the haplotypes of the selected relatives from the resulting phased data. We then input those haplotypes and HAPI2’s reconstructed parent haplotypes to TPBWT to detect IBD segments, again with phase correction disabled.

HAPI-RECAP leverages sex-specific genetic maps, and to analyze the SAMAFS data, we used the maps that Bhérer et al. produced ^19^, which are on GRCh37 coordinates. The 23andMe data we analyzed use GRCh38 positions, and to analyze those families, we used internally generated sex-specific maps on that genome build (produced using the methodology of Campbell et al. ^18^).

We compared the reconstructed genotypes with those produced by running HAPI2 on the families while including both parents and with (default) error options of --err_min 2 and --err_dist 5.

## Results

We ran HAPI2 on SNP array data from the SAMAFS and 23andMe nuclear families and then ran HAPI-RECAP using the IBD segments the reconstructed parents share with the siblings’ relatives. While each family includes data for both parents and between 3-12 children, we censored one or both parents’ data when performing these analyses. Below, we evaluate the reconstruction results using two primary metrics: the amount of DNA reconstructed and per-site error rate (Methods).

### Reconstructing genotypes for one parent

When data for siblings and one of their parents are available, HAPI2 reconstructs large fractions of the missing parent’s DNA. We ran it on the SAMAFS and 23andMe families (*n* = 114 and 220, respectively), omitting data in turn for the father and mother, with a total of 668 parents reconstructed. The amount of DNA HAPI2 reconstructs ranges from 77.6% to 99.97% (Figure 3), with the percentage varying with the number of children included. For families with three or four children, HAPI2 reconstructs 90.3% of the missing parent’s genotypes on average (median 91.8%), with a range of 77.6% to 96.2% (Figure 3, top right). Analyzing more children increases the amount of reconstructed DNA because a parent transmits both haplotypes across more of their genome to larger numbers of children. For families with seven or more children, HAPI2 reconstructs at least 95.9% of the parent’s DNA and up to 99.97%, with an average of 99.3% (median 99.6%) reconstructed (Figure 3, bottom right).

**Figure 3:**
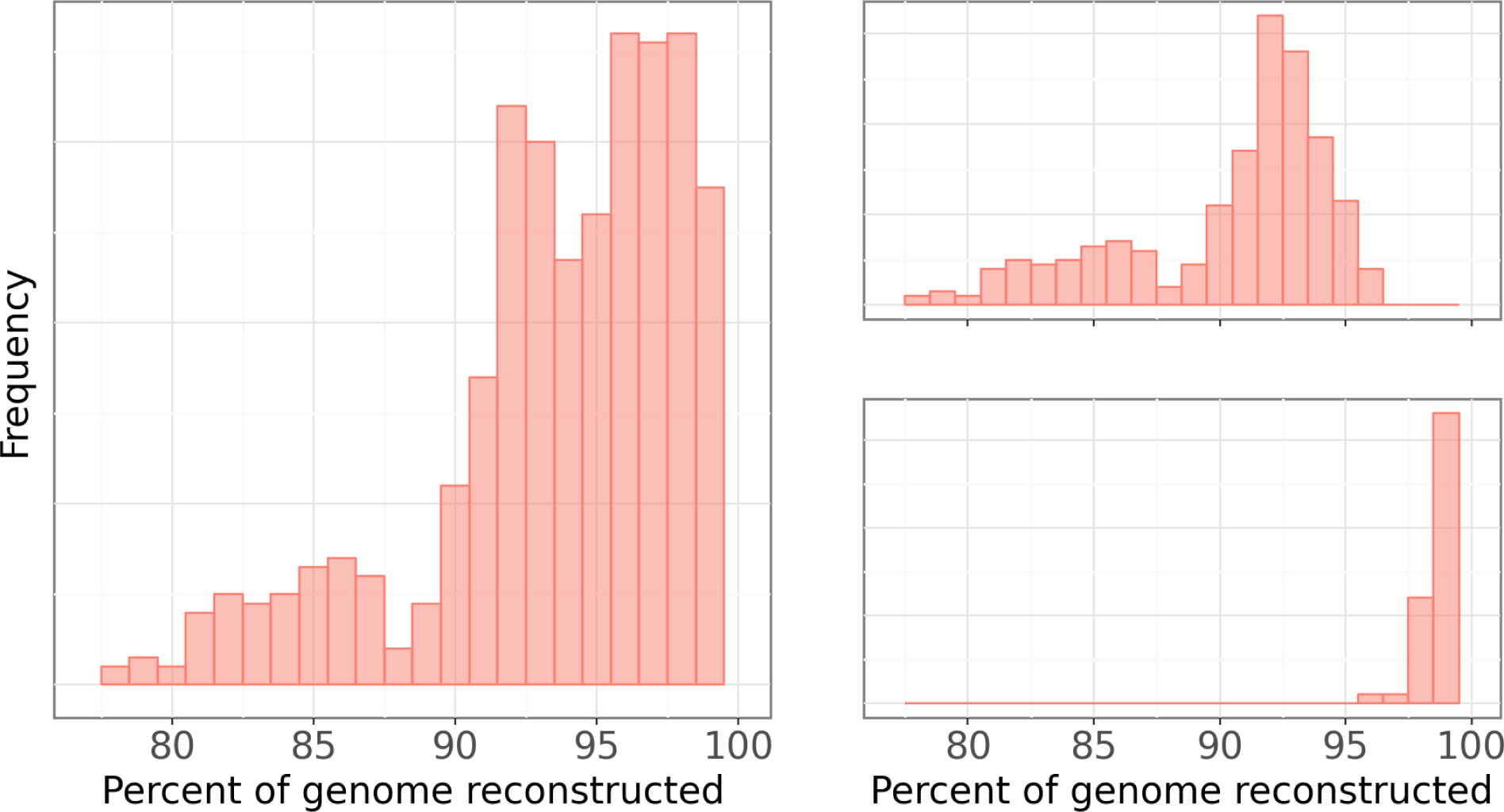
Histogram of the amount of DNA HAPI2 reconstructs when independently reconstructing the father and mother of each family. Figure depicts percentages from each parent in all families (left), in families with three or four children (top right), and in those with seven or more children (bottom right).

Crucially, the error rate in the reconstructed genotypes is low at < 10^−4^ per site on average. This is on par with or lower than the error rate of directly genotyping an individual ^33^.

### Reconstructing genotypes for both parents using only siblings and sex-specific genetic maps

When applied to many siblings, HAPI2’s reconstructed segments are generally long (the frequency of indistinguishable parent transmission decreases with increased numbers of siblings—see A.5: Distinguishing parent genotypes in the Appendix). Because of this and the large number of children, these segments also include many crossover events. HAPI-RECAP’s power to assign segments to the parents using sex-specific genetic maps is a function of the number of crossovers in a segment, so parent reconstruction without any IBD sharing is feasible with many siblings. We analyzed its performance on families with seven or more children. To be conservative, HAPI-RECAP only assigns a segment to the parents if |*LOD* | > 3. The method placed segments in all 35 families with ≥ 7 children (70 parents), reconstructing between 48.2% and 94.5% of each parent’s DNA, with an average of 77.7% and median of 82.1% reconstructed. The reconstruction quality remains high, with per-site error rates < 10^−4^.

### Reconstructing genotypes for both parents using siblings and their relatives

As described in Methods, HAPI-RECAP leverages the IBD segments detected between HAPI2’s reconstructed parents and the siblings’ relatives to resolve which parent transmitted different reconstructed segments. Of the 294 families with four or more children, 258 have at least one relative in either the SAMAFS or 23andMe cohorts that HAPI-RECAP could use to reconstruct the parents. As shown in Figure 4, HAPI-RECAP reconstructs 33.2% to 96.6% of the genotypes of each parent, and the amount varies with the number of siblings, as in the one-parent reconstruction analysis. For instance, in four-child families, the method reconstructs 33.2% to 83.2% of the parents’ DNA, with an average of 70.6% and a median of 72.0% reconstructed (Figure 4, top right). Families with seven or more children yield even more parent DNA, with a range of 70.7% to 96.6% reconstructed, and an average of 89.7% and median of 91.9% (Figure 4, bottom right).

**Figure 4:**
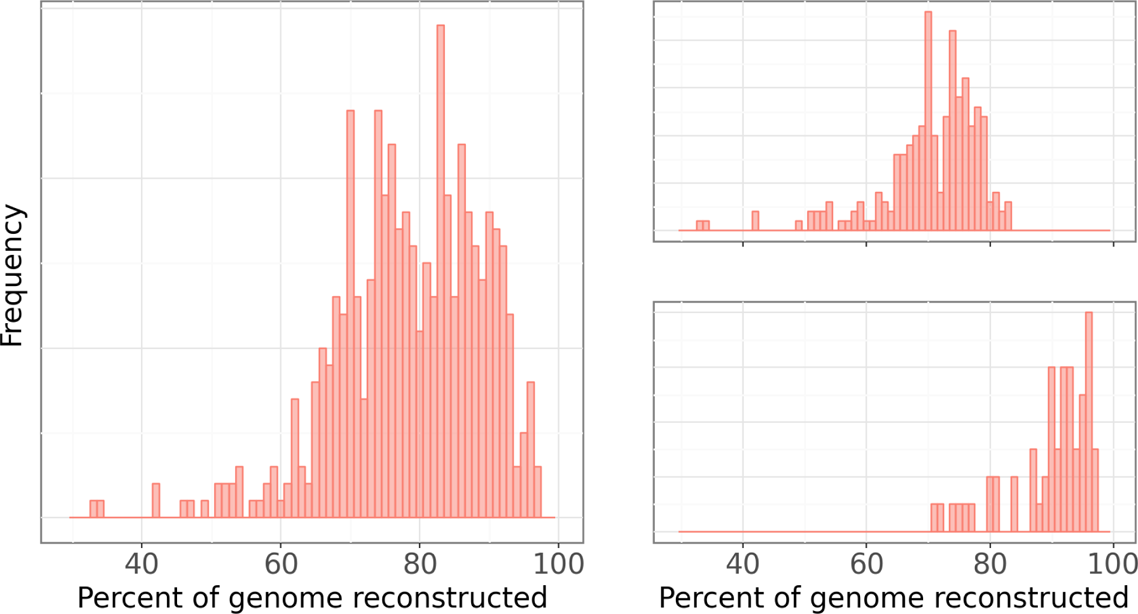
Histogram of the amount of DNA HAPI-RECAP jointly reconstructs for the father and mother of each family. Figure depicts percentages from each parent in all families (left), in families with four children (top right), and in those with seven or more children (bottom right).

HAPI-RECAP’s mean error rate for two-parent reconstruction is again very low at 1.4×10^−4^ per site (median 10^−4^), again meeting or exceeding the genotype quality of SNP arrays ^33^.

### Inferring parent sex and increasing reconstruction rate using sex-specific genetic maps

The analysis in the previous subsection considered only the autosomes and did not infer the sexes of the reconstructed parents, yet rich information for detecting these sexes is encoded in the locations of the crossovers the children inherited ^24^. HAPI-RECAP uses these crossovers to very reliably infer the parent sexes. This works by analyzing either a single reconstructed segment or a collection of these segments that the method assigns to the two unsexed parents using IBD.

Analysis of crossovers transmitted in multiple linked segments enables powerful and precise parent sex inference: HAPI-RECAP correctly infers the parents’ sexes in all 258 families whose parents it reconstructed using IBD segments (no errors). The statistical evidence for this inference is strong, with LOD scores ranging from 18.3-211 (mean 71.1).

Because crossovers within a single reconstructed segment also provide information about the parent sexes, it is possible to increase the amount of DNA reconstructed by inferring the sexes of the parents in regions that lack any IBD coverage. Placing segments with |*LOD* | > 3 adds an average of 4.2% of parent DNA (range 0.0-16.7%) to 69 families. However, the error rates in these segments is much higher at 9.5 × 10^−3^ per site or nearly 1%. Filtering to those families in which the method adds at least 10% of DNA to one parent improves the error rate to 10^−4^ per site and adds 10.3-16.7% of DNA to the parents (average 13.6%; median 14.1%) in 8 families.

Some families (*n* = 36) do not have relatives that meet HAPI-RECAP’s filters. In these cases, the application of sex-specific genetic maps enables parent reconstruction of each segment with |*LOD* | > 3. Such a strict threshold makes the reconstructed amounts highly variable, ranging between 2.4-88.2% in 34 families, but the quality remains high, with an error rate of 1.0 × 10^−3^. Limiting to families with at least one parent having 10% of their genome reconstructed leads to an error rate of < 10^−4^ and parent reconstruction rates ranging from 9.5-88.2% (43.0% mean and 40.7% median) in 27 families.

## Discussion

All of us inherit DNA from our ancestors and this forms the substrate for ancestor DNA reconstruction. The key problems in this reconstruction are (1) inferring a pedigree that includes the ancestors, (2) phasing the descendants, and (3) determining which haplotype segments descend from one or more ancestors. Reconstructing a nuclear family’s pedigree is very reliable ^14^, and all of the children’s DNA comes from their two parents—facts that aid in accurately reconstructing large fractions of the parents’ genomes.

Our approach applies family-based phasing ^17,20^, which is the gold standard ^34^ but is nevertheless subject to some ambiguities. For the problem of parent DNA reconstruction from siblings alone, a particularly impactful ambiguity arises when the two parents’ haplotype transmission patterns are indistinguishable (see A.5: Distinguishing parent genotypes in the Appendix). This not only affects the reconstruction at a single marker but, more importantly, prevents HAPI2 from determining whether the parents’ DNA was mixed from one side of the ambiguous region to the other.

Having ambiguous source parents for reconstructed segments on the same chromosome parallels the fundamentally ambiguous linkage of the parents’ DNA on different chromosomes. A closely related problem is linking the phased haplotypes for each chromosome of one person back to the source parents, or inter-chromosomal phasing. Recent work has addressed this problem using IBD segments a focal sample shares with their relatives ^21–23^: if the relative is connected through only one parent, the IBD segments will reside on haplotypes that parent transmitted. HAPI-RECAP applies this same IBD segment-based linkage to the reconstructed parent DNA. A key difference from inter-chromosomal phasing is that analyzing multiple siblings provides diploid chromosome reconstruction for the two parents in regions in which they transmitted both haplotypes (see A.6: Inferring that a parent transmitted only one haplotype).

Importantly, analyzing multiple children also enables the detection of crossovers the parents transmitted. Male and female genetic maps differ markedly, with males transmitting many of their crossovers near the telomeres and females transmitting overall more crossovers than males, with a more even distribution in placement throughout the length of a chromosome ^19^. We used these differences to infer the sexes of the reconstructed parents and, moreover, to place a segment on the parents even if no IBD segment overlaps it. Prior work showed that autosomal crossovers are highly informative for distinguishing paternal and maternal half-siblings and maternal and paternal grandparents ^24^, and here we found that even the crossovers transmitted within one reconstructed segment can provide sufficient information to determine the parents’ sexes.

A further source of information HAPI-RECAP does not use is that of statistical phase ^34^, which could help resolve ambiguities from family-based phasing. Errors in the siblings’ statistical phase and the high sensitivity of parent DNA reconstruction complicate the use of this information. A reconstruction error at a single marker can confound the DNA of the two parents, which often leads to confounded source parents across several megabases and results in numerous errors throughout a reconstructed segment.

Overall, HAPI-RECAP provides high quality reconstructed parent DNA. Its use of family-based phasing, IBD segments, and sex-specific genetic maps combine to enable precise inference of the DNA carried by the mother and father of a set of siblings. This and other related papers ^1,6,7,13^ demonstrate the potential for ancestor DNA reconstruction, a problem whose applications are vast. Moreover, the full promise of parent and general ancestor DNA reconstruction remains to be delivered: the completeness and accuracy of these reconstructions will continue increase as the sizes of both commercial and non-commercial genetic datasets continue to grow.

### A The HAPI2 algorithm

#### A.1 Detecting marker type

Like HAPI, HAPI2’s marker types are based on the parents’ genotypes and include: heterozygous for both parents, which are termed either *partly informative* and abbreviated PI for biallelic markers, or *fully informative for both parents* if the parents have distinct genotypes ^20^; heterozygous for one parent, which are called *fully informative for one parent* ^20^ and abbreviated FI1; and both parents homozygous, referred to as *uninformative* ^20^, abbreviated UN. The input format that HAPI2 uses (PLINK binary) allows only biallelic markers, so fully informative for both parent markers do not occur. However, it is possible to encode these multi-allelic variants as several biallelic markers with the same physical position.

Markers in which all individuals (children and any parents) are heterozygous have ambiguous phase, and HAPI2 omits them from its analysis. It also detects sites with Mendelian errors and does not phase them.

As used here, a site is *informative* if it provides information about which haplotypes a parent transmitted to the children, so a site is informative for any heterozygous parent and uninformative if both parents are homozygous. HAPI2 generates states in its HMM only at informative sites: all others either do not require phase (all individuals are homozygous) or, if the parents are homozygous for opposite genotypes, the heterozygous children are trivial to phase.

Table 1 lists all possible genotypes for a set of children and the parent genotypes consistent with these when one or both parents are ungenotyped. Table 2 gives the marker types when the parents’ data are given. HAPI2 uses applies this to infer marker types.

**Table 1:**
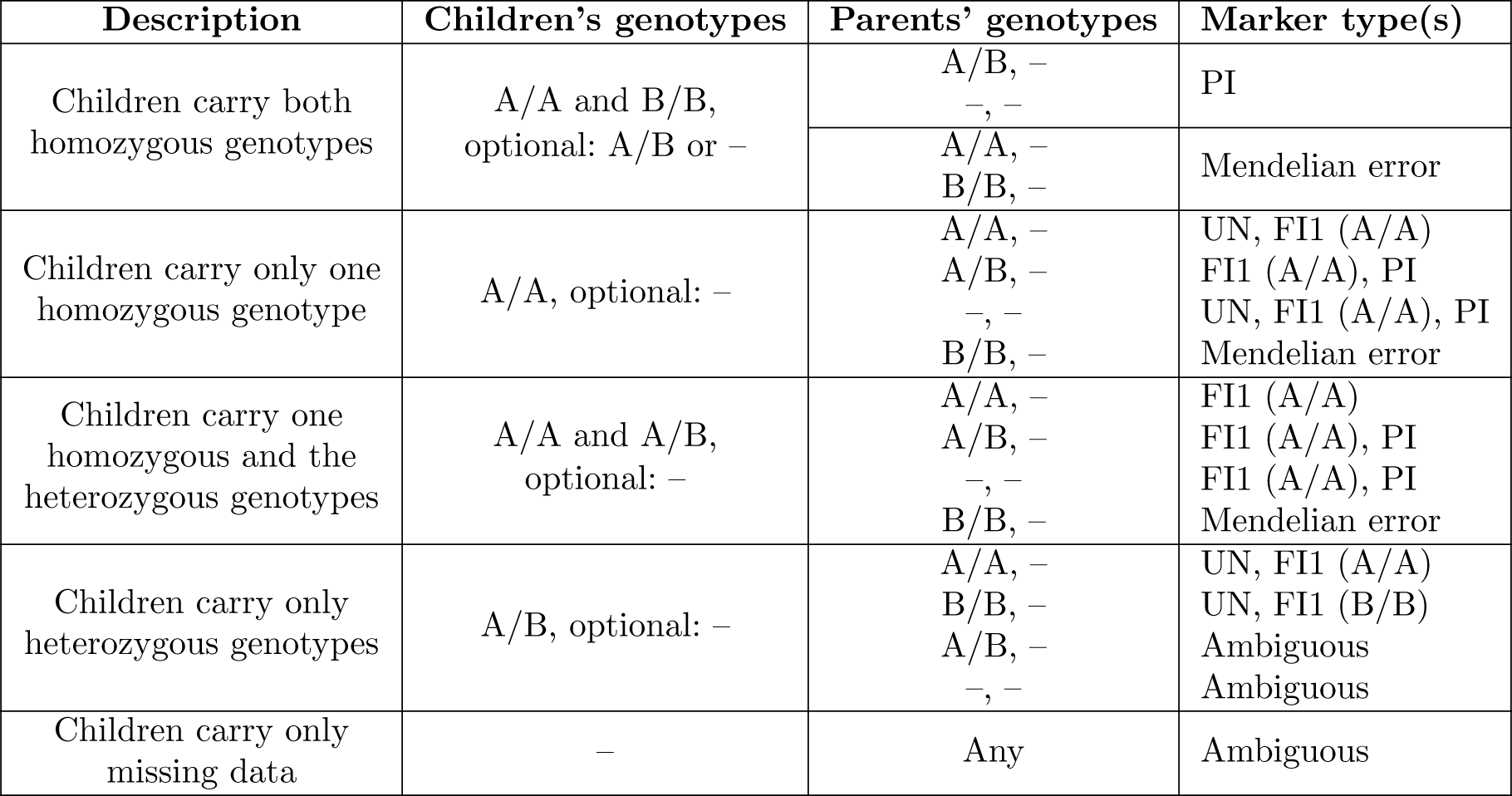
Marker types HAPI2 uses on the autosomes for the given children’s genotypes and non-missing genotypes for one or neither parent. Some combinations of genotypes have Mendelian errors or yield ambiguous phase, as indicated; – represents missing data. For FI1 markers, the table lists in parentheses the implied or given genotype for the homozygous parent, e.g., FI1 (A/A).

**Table 2:** Marker types HAPI2 uses on the autosomes given data for both parents. Some combinations of genotypes have Mendelian errors, as indicated; if all non-missing genotypes are A/B, the phase is ambiguous (last row); – represents missing data. For FI1 markers, the table lists in parentheses the given genotype for the homozygous parent, e.g., FI1 (A/A).

| Description | Parents' genotypes | Children's genotypes | Marker type |
| --- | --- | --- | --- |
| Parents both homozygous | A/A, A/A | A/A or –<br>Any A/B or B/B | UN<br>Mendelian error |
|  | A/A, B/B | A/B or –<br>Any A/A or B/B | UN<br>Mendelian error |
| One parent heterozygous | A/B, A/A | A/B or A/A or –<br>Any B/B | FI1 (A/A)<br>Mendelian error |
| Both parents heterozygous | A/B, A/B | A/B or A/A or B/B or –<br>All A/B or – | PI<br>Ambiguous |

Sites that are potentially uninformative include those where all children are homozygous for the same allele (e.g., all are A/A) and those where all children are heterozygous and the genotypes of any genotyped parents are homozygous. While the missing parents for these sites *could* be heterozygous, in many cases the parents are homozygous and the site is truly uninformative. The only way a missing parent’s genotype could be heterozygous at such sites is if the parent transmitted only one haplotype (since all children inherited the same allele in these cases). HAPI2 detects this, as described below (see Inferring that a parent transmitted only one haplotype), but when not applying that methodology, it treats the site as uninformative, assigns homozygous genotypes to both parents, phases the children if they are heterozygous (and therefore the parents must be homozygous for different alleles), and proceeds to analyzing the next marker.

#### A.2 Generating partial states

Given the set of all possible parent genotypes at a site, HAPI2 constructs partial states. Partial states correspond to only one way of phasing each heterozygous parent and include incompletely defined inheritance vectors (IVs) for the children. (Later, full states include fully defined IVs and incorporate all possible parent phase assignments.)

If the marker can be FI1 and both parents are missing data, HAPI2 constructs two partial states, one for each possible parent genotype (e.g., parent 1 A/B and parent 2 A/A; and parent 1 A/A and parent 2 A/B). Further, if the marker can be PI, the method constructs a partial state with this genotype assignment to the parents (e.g., both parent 1 and parent 2 A/B). Thus there are at least one and no more than three partial states at each marker when one or both parents are missing. If neither parent is missing, HAPI2 only constructs one partial state as dictated by the parents’ genotypes (Table 2).

Partial states that are FI1 enable the definition of half of the IV—the vector elements corresponding to the heterozygous parent’s transmitted haplotype. For example, if parent 1 is A/A and parent 2 is heterozygous A/B with the A and B alleles assigned to haplotypes 0 and 1 respectively, children with genotype A/A must have inherited haplotype 0 from parent 2 and those with genotype A/B inherited haplotype 1 (Table 3). In the partial state, the IV elements corresponding to the homozygous parent are undefined, and if a child is missing data, both their corresponding IV elements are undefined.

**Table 3:** Example partial states of FI1 and PI types. The parent haplotypes are in VCF notation, with parent 1 on the left and 2 on the right. Genotypes for four children are listed in the same order as in the IVs (e.g., the first child’s IV values are the first two IV elements). Partial FI1 states indicate which haplotype the heterozygous parent transmitted to the children. A homozygous parent (in this example, parent 1) provides no information about which haplotype they transmitted, so the partial state lists undefined values for that parent’s transmissions in the IV, indicated here as −. PI states provide fully defined IV values for homozygous children and ambiguous IV values for heterozygous children, indicated with a(see Ambiguous inheritance vector values). A child with missing data also provides no information about which haplotypes they received, so their IV values are undefined.

| Marker type | Parents' haplotypes | Children's genotypes |  |  |  | IV |  |  |  |
| --- | --- | --- | --- | --- | --- | --- | --- | --- | --- |
| FI1 | A\|A A\|B | A/B | A/A | A/A | A/B | -1 | -0 | -0 | -1 |
|  |  | A/B | A/A | A/A | – | -1 | -0 | -0 | -- |
| PI | A\|B A\|B | B/B | A/B | – | A/A | 11 | 01? | -- | 00 |

Partial states that are PI enable a complete definition of IV elements for all homozygous children but an incomplete—ambiguous—IV assignment for all heterozygous children. For example, assuming both parents have their A and B alleles assigned to haplotypes 0 and 1, respectively, children that are homozygous A/A each inherited haplotype 0 from each parent, so their IV values are 00; in turn, the IV values for all children with B/B genotypes are 11. A/B heterozygous children could have IV values of either 01 or 10, depending on their phase, and the partial state adopts the original HAPI’s encoding of *ambiguous* IV values ^20^, which we describe in the next subsection (Table 3). Lastly, as in the FI1 case, the IV elements for a child with missing data are undefined.

#### A.3 Ambiguous inheritance vector values

Ambiguous IV values apply only at PI states and only to children that are heterozygous. In these cases, the child could have inherited either allele from each parent, so either of the two possible phases may be correct (e.g., using VCF format notation, the child’s phase could be A|B or B|A). Considering only data at the current marker, the phase is completely ambiguous. An ambiguous IV value corresponds to two IV values for the child, and these are exactly inverted from one another, e.g., 01 and 10 as in the previous subsection. The reason for this is that swapping the child’s phase corresponds to inverting which allele each parent transmitted, and this inverts their transmitted haplotypes (the parents are both heterozygous). There are four possible (unambiguous) IV values for each child and two ambiguous IV values: (01 or 10) and (00 or 11). To encode an ambiguous IV value, HAPI2 stores one of the valid IVs (say 01) and then stores a separate flag indicating the ambiguity. In this paper, we denote the ambiguity by listing one of the valid IVs followed by acharacter (Table 3).

As detailed previously ^20^, use of ambiguous IV values avoids a state space explosion since otherwise there would be 2*^h^* IVs for each of the four parent phase assignments in PI states, where *h* is the number of heterozygous children at a site.

#### A.4 Generating full states

At each informative marker, HAPI2 builds full states from each partial state and incorporates these into its HMM, transitioning to them from states at the previous informative site. Initially, at the first informative site (regardless of type) on a chromosome, no previous states exist and so there is no information to specify undefined and/or ambiguous IV values. As such, HAPI2 merely copies the partial states (including any undefined or ambiguous IV values) to generate the first set of full states, and it assigns each of these a total recombination count of 0. HAPI2 adds defined values and resolves ambiguous IV values during back tracing. The phase assignments of the parents are arbitrary in partial states and it suffices to retain these arbitrary phase assignment for the first site that is heterozygous in each parent: HAPI2 phases downstream sites relative to this. When both parents are missing data, for the first informative site on an autosome that may be FI1, incorporating only FI1 states that are heterozygous for parent 1 and not parent 2 suffices. This is because we do not have sex information or, at this stage (i.e., absent IBD or sex-specific map-based information), the ability to link parent 1 on chromosome 1 to parent 1 or 2 on other chromosomes. Therefore, the code arbitrarily assigns a label (parent 1) to the first heterozygous parent, allowing downstream methods to determine which reconstructed segments belong to the same parent. An alternative is to include states for both possibilities. This would yield equivalent state paths with opposite parent genotype assignments, but no additional information and, indeed, would require additional computation. As such, HAPI2 only includes states with parent 1 assigned heterozygous at this first site.

Given states at the previous informative marker, HAPI2 generates full states by looping over the previous states and calculating the IV values that result from transitioning from each one to each partial state, building a full state for every combination. We describe this process initially for full states with the same parent phase as in the partial state and then discuss generating full states with alternate phase assignments below. In a full state with parent phase matching the partial state, HAPI2 copies the defined IV elements from the partial state—these transmissions are implied by the parents’ phase and the children’s genotypes, and the previous state being transitioned from does not alter this. For any IV values that are undefined in the partial state (including for a child with missing data), HAPI2 copies the IV values from the previous state. This is consistent with minimum-recombinant phasing: undefined IV elements occur when no haplotype transmission information is available, and it suffices to avoid introducing any recombinations that have no data to support them. If an IV element is undefined in the previous state, it remains undefined in the full state being generated, so for example, a series of states that are heterozygous only for parent 1 at the beginning of the chromosome retain undefined IV elements corresponding to parent 2’s transmissions. The method defines these IV elements at the first state that is informative for parent 2.

For ambiguous IV values in the partial state, several possibilities exist. (1) The previous state’s IV may be unambiguous and non-recombinant relative to one of the possible IVs in the partial state (e.g., the previous state’s IV is 01 and the partial state’s ambiguous IV is (01 or 10)). In this case, HAPI2 uses the same (unambiguous) IV as in the previous state, consistent with minimum recombinant phasing; this is quite likely to be correct since the alternate IV would induce two recombinations—one on each haplotype transmitted to this child—relative to the previous state. (2) The previous state’s IV is unambiguous but requires a recombination to transition to either of the IVs encoded in the partial state (e.g., the previous state’s IV is 11 and the partial state’s IV is (01 or 10)). As the child could be phased in either orientation, and as each possible IV implies exactly one recombination, HAPI2 assigns an ambiguous IV to the full state in this case. This ambiguous IV gets resolved in the same way as in HAPI ^20^; see Back tracing. (3) The previous state’s IV is ambiguous. In this case, HAPI2 assigns the ambiguous IV from the partial state to the full state; notably, this IV may differ from and therefore be recombinant relative to that of the previous state.

It is straightforward to deduce the IV corresponding to each possible parent phase from the partial state. In FI1 partial states, inverting the heterozygous parent’s phase merely inverts the corresponding IV elements: the same parent still transmitted the same alleles to the children, but the haplotype those alleles derive from is inverted. This principle also applies to homozygous children in PI partial states: if, e.g., parent 1’s phase is inverted, all IV values corresponding to that parent’s haplotype transmissions must be inverted. For heterozygous children in PI partial states, if both parents’ phases are inverted, this maps to the same ambiguous IV values in the children, and HAPI2 does not change their ambiguous IVs. If one parent’s phase is inverted, the ambiguous IVs must change, e.g., from (01 or 10) to (00 or 11). HAPI2 applies these adjustments to the IVs as it generates full states corresponding to all possible parent phases for each partial state.

Since an FI1 state has only two parent phasings and a PI state has four (two for each parent), the total number of states generated at each marker is (2*n_FI1_* + 4*n_PI_*) · *p*, where *n_FI1_ ∈ {*0, 1, 2} is the number of FI1 partial states, *n_PI_ ∈ {*0, 1} is the number of PI states, and *p* is the number of states at the previous informative marker. This increase relative to *p* could lead to a massive state space after several markers, but some of these states have the same or equivalent (see Optimizations) IV and therefore are only represented by a single state in the HMM. That is, HAPI2 stores states with equivalent IVs—the only component that affects downstream processing—in the same state, while indicating in the state whether multiple different parent phases and/or parent genotypes yield the same number of recombinations. Each state stores the minimum number of recombinations required to transition to it and stores the previous state(s) that yield this minimum. Furthermore, HAPI2 deletes any states that are guaranteed to be suboptimal (see Optimizations). Given this, it remains efficient even when analyzing families with many children.

Calculating the number of recombinations relative to a previous state is straightforward: for any defined (including ambiguous) IVs, the Hamming distance gives the recombination count. HAPI2 stores the minimum total recombinations required to transition to each state and thus stores the sum of the Hamming distance and the minimum recombination count in the previous state. However, as this is a minimum recombination count, if a different previous state yields fewer recombinations to transition to some state, HAPI2 retains the minimum count and only stores the previous states that yield this count.

#### A.5 Distinguishing parent genotypes

When both parents’ data are missing, the approach HAPI2 uses to distinguish parent 1 and parent 2’s genotypes is a function of the children’s genotypes (the only observed data), and these are a reflection of the latent haplotype transmissions from each parent—which HAPI2 encodes in IVs. At FI1 markers, if the two parent’s haplotype transmission patterns are distinct, it is straightforward for HAPI2 to detect when parent 1 is heterozygous and when parent 2 is. For example, if the IV at a marker in a four child family is 01 10 11 00 (i.e., the children inherited each of the four possible combinations of haplotypes) then if parent 1 is A/B with haplotypes 0 and 1 carrying the A and B alleles, respectively, and if parent 2 is A/A, the children’s genotypes would be A/A, B/A, B/A, and A/A. This differs from the genotypes if instead parent 2 is A/B with haplotypes 0 and 1 carrying the A and B alleles, respectively, with parent 1 being A/A, which corresponds to child genotypes of A/B, A/A, A/B, and A/A: two of the four children’s genotypes are distinct (child 1 and 2).

In cases where the IVs of the two parents are indistinguishable, HAPI2 cannot determine which parent is heterozygous at an FI1 marker. For example, if a marker’s IV in a four child family is 00 11 00 11, then regardless of whether parent 1 or parent 2 is A/B, assuming the A and B alleles are on haplotypes 0 and 1, respectively, and if the homozygous parent is A/A, the children’s genotypes would be A/A, A/B, A/A, and A/B.

The two parents’ IVs need not match perfectly for their haplotype transmissions to be indistinguishable. They can be either the same or the inverse of each other (e.g., 01 10 01 10) and still produce the same observed data: the inverted IV corresponds to inverted haplotype assignments, which, as noted earlier, is chosen arbitrarily at the beginning of the chromosome. Thus, in general, of the 2^2c^ possible IVs, 2*^c^*^+1^ correspond to indistinguishable parent haplotype transmissions: any IV for parent *i*, of which there are 2*^c^*, has one identical and one inverted IV for the other parent, yielding 2 *·* 2*^c^* = 2*^c^*^+1^ IVs. This means that the fraction of IVs in which the two parents are indistinguishable depends on *c* and is considerably lower in large families (2*^c^*^+1^*/*2^2c^ decreases as *c* increases).

Interestingly, markers that are necessarily PI and not FI1—i.e., where the children carry both homozygous genotypes (Table 1)—can still be phased in these regions. The children’s homozygous genotypes coupled with the IV from previous states imply the parents’ phase. In fact, these PI markers are those where HAPI2 initially detects that the two parents’ haplotype transmissions have become indistinguishable.

HAPI2 detects when a state has IVs that make the parents indistinguishable at FI1 markers. In these cases, it only produces states that are heterozygous for parent 1 and marks the site as parent-genotype ambiguous. A further consequence of the parents being indistinguishable is that the reconstructed genotypes upstream and downstream of such regions cannot be linked to one another. Just as HAPI2 does not infer the source parent of the genotypes on distinct chromosomes, the unlinked nature of reconstructed genotypes on either side of these parent ambiguities mean that other sources of information are needed to assign the segments to the same parent. Both IBD segments and, with sufficiently informative crossover events, sex-specific genetic maps can link the reconstructed DNA to the same parent.

#### A.6 Inferring that a parent transmitted only one haplotype

During processing, in most cases, when a child inherited a recombined haplotype relative to a previous site, evidence of this is simple to detect; for example, the IVs in even the partial states differ from those of previous markers. However, if a parent with missing data transmits only one haplotype to all of the children, those children’s genotypes will be consistent with that parent being homozygous even at sites where they are truly heterozygous. Absent some means of detecting this case, HAPI2, which seeks minimum-recombinant phase, would retain the same IV as at previous sites: the data can be phased without introducing this recombination. (The recombination may manifest later at a site with evidence of heterozygosity in the parent, i.e., once an additional recombination occurs to an IV with both of the parent’s haplotypes transmitted.)

To detect probable locations where a parent transmitted only one haplotype, in which case HAPI2 will reconstruct only that haplotype for the parent, the algorithm adds a artificial FI1 states in certain circumstances. Specifically, after encountering a pre-specified number of sites that are all uninformative for a given parent (termed *F_i_* later; by default 150 markers), if the IV has ≤ 2 children inheriting the minority haplotype (so 1 or 2 recombinations are needed for the parent to have transmitted a single haplotype), HAPI2 adds these forced informative states. The effect of this is to introduce an IV where all children received the same haplotype from the parent (since they will have all inherited the same allele). The first such site will necessarily be recombined relative to previous sites (as noted in the last paragraph). HAPI2 inserts these forced FI1 states at regular (user-specified) intervals (by default every 50 markers) until it encounters a site detected as heterozygous in the parent.

In some cases, the parent truly has a run of homozygosity (ROH) ^35^ and ideally HAPI2 would reconstruct the parent as homozygous. If the interval is sufficiently short, HAPI2’s error detection (below) can filter even these forced FI1 states and, in that case, would reconstruct the region as an ROH. In other cases, HAPI2 will infer a half-missing genotype for the parent, which may not be correct, but is conservative.

When scoring HAPI2’s accuracy, we count a half-missing genotype as correct if the single reconstructed allele matches one of the parent’s alleles, and we count a full homozygous genotype as incorrect if the parent is heterozygous (see Validation and quality metrics for reconstructed genotypes).

It is possible for both parents to transmit a single haplotype in the same region, and the above procedure applies in this case, with HAPI2 introducing the forced FI1 markers independently for each parent.

The back tracing procedure infers half-missing genotypes for a parent conservatively: it assigns them at all sites where either the up- or downstream flanking informative marker has an IV where a parent transmitted only one haplotype. This means that even though the first forced FI1 state occurs at least 150 markers (by default) from an informative site, the uninformative sites preceding the first and following the last forced FI1 states are reconstructed with just one haplotype.

#### A.7 X chromosome phasing

HAPI2 supports phasing of the X chromosome, which, after early processing, executes the same code as for autosomes. The process begins by checking for males called heterozygous, which the method recodes as missing, and then detecting the marker type of a site. Marker type detection on the X chromosome is unique because the hemizygosity of males alters the observable genotypes and reasoning about missing parent data. Table 4 lists the possible parent and child genotypes when one or both parents are missing data and gives the inferred marker types in each case on the X chromosome (note that the PI marker type is impossible on the X chromosome since the father is never heterozygous). When the father’s genotype is missing, HAPI2 either imputes it by Mendelian inheritance rules using the daughters’ genotypes or, for FI1 states, generates a state for each of the two possible homozygous genotypes for the father. Likewise, the code also attempts to infer the mother’s genotype when it is missing (Table 4). One special marker type (indicated with a * in Table 4) is often uninformative but is sometimes nearly completely ambiguous. Specifically, if all sons inherited the same haplotype from the mother and all daughters (who are all heterozygous) inherited the other haplotype, the mother could be either homozygous or heterozygous, depending on the phase of the daughters. In this case, the only allele that can be inferred is the one the mother transmitted to the sons. HAPI2 flags these special marker types, treats them initially as uninformative, checks the IVs during back tracing, and assigns only one allele to the mother when needed.

**Table 4:** Marker types HAPI2 uses on the X chromosome for the given children’s genotypes and non-missing genotypes for one or neither parent. Some combinations of genotypes have Mendelian errors or yield ambiguous phase, as indicated; – represents missing data and we list male genotypes as if they are diploid homozygous. For FI1 markers, the table lists in parentheses the genotype of the father, e.g., FI1 (A/A), or, if this is not listed, the father’s genotype is the same as any homozygous daughters. When there are no homozygous daughters, the father could be either A/A or B/B, which the table indicates as FI1 (?), and HAPI2 creates states corresponding to both possibilities. One special case marker type is indicated with * and the parents’ genotypes may in fact be nearly completely ambiguous as described in the text.

| Parents' genotypes |  | Children's genotypes | Marker type(s) |
| --- | --- | --- | --- |
| Mother | Father |  |  |
| A/A | – | Any B/B<br>Both A/A and A/B daughters<br>None of the above | Mendelian error<br>Mendelian error<br>UN |
| A/B | – | Some A/A and B/B daughters<br>Any A/A or B/B daughter<br>Only A/B, –, or no daughters; some sons<br>Only A/B, –, or no daughters; – or no sons | Mendelian error<br>FI1 (see caption)<br>FI1 (?)<br>Ambiguous |
| – | A/A | Any B/B daughter<br>A/A and (B/B or A/B), optional: –<br>Neither of above | Mendelian error<br>FI1 (A/A)<br>UN, FI1 (A/A) |
| – | – | Some A/A and B/B daughters<br>A/A and (A/B or B/B), optional: –<br>Only A/B, –, or no daughters; some sons<br>Only A/B, –, or no daughters; – or no sons<br>None of the above | Mendelian error<br>FI1 (see caption)<br>UN, FI1 (?), *<br>Ambiguous<br>UN, FI1 (see caption) |

Partial state generation for X chromosome markers is straightforward and in fact, works by temporarily using diploid genotypes for the sons by joining their original hemizygous call with the father’s hemizygous call. Given these sons’ diplotized genotypes and the rest of the family’s data, the same code that generates partial states for the autosomes applies, though we use defined (and arbitrarily) IV values for the paternal transmissions. HAPI2 also detects regions where the mother transmitted only one haplotype on the X chromosome, similar to the autosomal case (see Inferring that a parent transmitted only one haplotype).

#### A.8 Detecting genotyping errors

Genotyping errors can confound family-based phasing by implying spurious recombinations that prematurely end a region in which a parent transmitted only one haplotype or that suggest that all children recombined in a short interval (e.g., if a truly homozygous parent is called heterozygous), among other issues. HAPI2 includes an option to detect markers where one or more family members may have an error and mark them as erroneous in its output. This operates through two command-line options: --err_dist #, which specifies how many informative markers in a row to potentially group as arising from a single error (referred to as *D* below), and --err_min #, which specifies the minimum number of recombinations a series of up to *D* informative sites must include in order to be called erroneous (referred to as *E* below; --err_min 0 disables error detection). For example, a site with one genotyping error in a child can appear as one recombination at that site followed by another recombination back to the previous haplotype at the next informative marker, so would produce two recombinations; using --err_min 2 would flag such a marker as an error.

HAPI2’s error detection works by considering transitions from states at informative markers upstream of the previous informative site: if such a transition yields a lower recombination count (including the penalties we include for incorporating the error, as described below), HAPI2 marks the intervening informative sites as errors during back tracing. Thus the error detection analysis considers only informative markers—those with states in the HMM—and we index these sequentially, e.g., as informative marker *m, m* + 1, and so on. This ignores any uninformative or ambiguous sites and those with Mendelian errors that may occur between informative markers. Although some uninformative and ambiguous sites may include genotyping errors, these do not impact HAPI2’s results other than at those sites and so have limited impact on the phasing results.

The algorithm proceeds by tracking a list *L_e_* of states upstream of the previous informative site to potentially transition from. Any state at a site that is truly informative (i.e., not the artificial informative states discussed in Inferring that a parent transmitted only one haplotype) is potentially eligible. Generally HAPI2 adds states to *L_e_*, but a state at informative marker *m* that transitions to informative marker *m* + 1 with zero recombinations does not need to be included: the state at informative marker *m* + 1 should be incorporated on any state path that the immediately previous state is on. One exception to this is that the method always includes states with undefined IV elements in *L_e_* because such states transition with zero recombinations even to a site with an error if the error(s) impact the undefined IV elements.

Given *L_e_*, HAPI2 follows the algorithm described in Generating full states to form transitions from those upstream states to the current marker. As in the no-error case, the method counts the number of recombinations that result from each transition and also adds a penalty “recombination” term equal to *E −* 0.5 + 0.1(*S −* 1), where *S ≤ D* is the number of markers that the transition would mark as an error (i.e., the number of informative markers that separate the current site from the state being transitioned from). For example, suppose marker *m* + 1 has an error that produces two recombinations (above) but the optimal state path has the same IVs at markers *m* and *m* + 2. Then if *E* = 2, the error detection code will consider a transition from the optimal states at *m* to *m* + 2, and this will produce 0 recombinations plus a penalty of *E −* 0.5 = 1.5 recombinations (this skips only one marker, so *S* = 1 and 0.1(*S −* 1) = 0). As 1.5 < 2, HAPI2 will assign this new, lower recombination count to the optimal state at marker *m* + 2 and update the previous state that transitions to it as the one at marker *m*.

In otherwise ambiguous cases where the number of recombinations (including the penalty term) that arise from a path with an error is equal to a path without an error, HAPI2 chooses the state path without an error.

As HAPI2 completes the analysis at informative marker *m*, it removes any state *s* from *L_e_* if *m − m_s_ > D*, where *m_s_* is the informative marker index of state *s*. These are too distant from informative marker *m*+1 to be considered at that next site. Further, it removes any state in *L_e_* that transitions with zero recombinations (ignoring penalties) to the current marker; again, it suffices to consider the state at the current marker downstream since any state path that includes the earlier state should also include the current state.

Importantly, errors can occur at the beginning of a chromosome, and in addition to considering transitions from previous states in *L_e_*, when the number of previous informative markers on a chromosome is ≤ *D*, HAPI2 incorporates states with no previous marker. These include the same penalty in their recombination count as noted above, and if the optimal state path includes this type of state, the method marks any omitted markers as errors during back tracing.

In some cases, including those where one or both parents transmitted only one haplotype, the informative marker density can be low, with many uninformative or ambiguous sites between informative markers. Because of this, HAPI2 only considers transitioning from a state *s* in *L_e_* if the distance from the current to that marker is *M − M_s_ ≤ F_i_* + 50, where *M* and *M_s_* are the marker indexes (counting all sites) of the current marker and that of state *s*, respectively, and *F_i_* is the number of sequential uninformative markers required before HAPI2 adds an artificial informative marker (see Inferring that a parent transmitted only one haplotype). This ensures that artificial informative markers get added. Despite this limit, HAPI2 always considers transitions from states at informative marker *m −* 2, where *m* is the current informative marker (i.e., the informative marker immediately before the previous informative marker). This marker count also applies at the beginning of the chromosome.

#### A.9 Optimizations

Like its predecessor, HAPI2 provides efficient family-based phasing. Three of its key optimizations are also part of HAPI ^20^: it uses ambiguous IV values (see Ambiguous inheritance vector values above), it only stores one state for all equivalent IVs, and it removes states that are guaranteed to be suboptimal. The implementation of the latter two optimizations is somewhat different in HAPI2 than in HAPI.

Any two IVs that have exactly inverted elements for either parent (or both) are equivalent to one another ^20^. Such IVs correspond only to different haplotype assignments in the parent(s), and any downstream sites are readily phased in either orientation given a choice at some marker. Given this, HAPI2 uses a canonical IV representation that all four equivalent IVs map to. This mapping incorporates ambiguous IV values (i.e., to be equivalent, the same children must be ambiguous) and requires the same set of defined elements (i.e., an IV that is undefined for a different set of children than another IV maps to a distinct canonical IV). During back tracing, HAPI2 updates the IVs so that the final output reflects the parent haplotype transmissions and inferred recombinations.

When generating full states at a marker, HAPI2 tracks and stores both the minimum and maximum recombination counts among these. Because downstream recombinations can lead to any IV from any other, it suffices to remove states with recombination counts that are greater than the number that would result from transitioning from the minimum-count state to its IV. Several ways for implementing this are possible, and, in fact, one could analyze the IVs among all states, not just the state with the minimum recombination count. For simplicity, HAPI2 removes a state *d* if *r_d_ ≥* 2*c* + *r_m_*, where *r_s_* is the total number of recombinations in a given state *s* and *m* is the minimum recombination state at the current marker. Thus even if it were necessary for *all* children in the minimum state to recombine downstream, that minimum state would provide the same or lower overall recombination count than the state to be removed.

HAPI2 stores all IVs using 64 bits, with each vector element stored in a single bit. Storing ambiguous IV values uses another 64 bits, as does representing which elements in an IV are defined. Bitwise operations provide an efficient and convenient means of manipulating and comparing IVs, and HAPI2 uses these operations.

#### A.10 Back tracing

After reaching the end of a chromosome, HAPI2 applies the phase defined by the minimum state path(s) in its HMM. This works by first identifying the state(s) at the last informative marker with minimum recombinations and then iteratively tracing back to the state(s) at each previous informative marker that transition to the current state(s). The algorithm tracks and indicates regions where the two parents are indistinguishable and reports if, e.g., there is ambiguity in whether one or both parents are heterozygous, if the parents’ phase is ambiguous, and if any IV elements are ambiguous (such as at uninformative sites between two markers that are recombined relative to each other). Although the HMM-generation code seeks to avoid introducing states that are equivalent but with opposite parents assigned as heterozygous, the back tracing code makes a final check for this and, if it encounters such a state path, only retains one of the possibilities while indicating at the beginning of such a region that the chosen heterozygous parent is arbitrary (i.e., that the two parents are indistinguishable at that first marker). The algorithm traces back all minimum recombinant states; however, if, due to errors, one state traces back to a different marker than another, HAPI2 arbitrarily chooses a state path to continue tracing and stops tracing back state paths that lead to a different marker.

During back tracing, HAPI2 resolves ambiguous IV values (see Ambiguous inheritance vector values) and propagates IV elements back to positions with undefined values. For ambiguous IV values, typically down-stream fully informative sites can be phased consistent with one of the two possible IVs, and HAPI2 resolves the ambiguity to match that later IV. In rare cases (e.g., a non-crossover gene conversion that yields a single SNP recombination that reverts to the original haplotype ^29^), which parent transmitted the recombinant genotype remains ambiguous, and HAPI2 represents this ambiguity in its final phasing result.

The code assigns only one allele to a parent in regions they were likely to have transmitted only one haplotype (see Inferring that a parent transmitted only one haplotype). It also assigns only one allele to a parent at uninformative sites at the beginning (respectively, end) of a chromosome if the IV for that parent at the first (last) informative marker has only one child inheriting one of the haplotypes.

## Declaration of interests

E.M.J., K.F.M, W.A.F., and A.L.W. are current or former employees of and may hold equity interest in 23andMe, Inc. Y.Q. was an intern at 23andMe, Inc. for part of her work on this paper. A.L.W. is the owner of HAPI-DNA LLC. All other authors declare no competing interests.

## Data and code availability

The HAPI-RECAP and HAPI2 code bases will be made available upon publication. Individuals with direct-to-consumer company genetic data for three or more siblings and one parent can reconstruct the other parent’s DNA for free at https://hapi-dna.org/hapi/. This web-based tool executes on the user’s computer so does not transfer data to HAPI-DNA.

## Acknowledgments

We thank the research participants who tested with 23andMe and members of the SAMAFS for making this work possible. We also thank Adam Auton for generating the GRCh38 sex-specific genetic maps. Funding for this work was provided by NIH grants R35 GM133805, U54 HG013247, and R01 MD012564 and by 23andMe, Inc. Some of the computing was performed on a cluster administered by the Biotechnology Resource Center at Cornell University.

The following members of the 23andMe Research Team contributed to this study:

Stella Aslibekyan, Adam Auton, Elizabeth Babalola, Robert K. Bell, Jessica Bielenberg, Ninad S. Chaudhary, Zayn Cochinwala, Sayantan Das, Emily DelloRusso, Payam Dibaeinia, Sarah L. Elson, Nicholas Eriksson, Chris Eijsbouts, Teresa Filshtein, Pierre Fontanillas, Davide Foletti, Will Freyman, Zach Fuller, Julie M. Granka, Chris German, E^á^daoin Harney, Alejandro Hernandez, Barry Hicks, David A. Hinds, M. Reza Jabalameli, Ethan M. Jewett, Yunxuan Jiang, Sotiris Karagounis, Lucy Kaufmann, Matt Kmiecik, Katelyn Kukar, Alan Kwong, Keng-Han Lin, Yanyu Liang, Bianca A. Llamas, Aly Khan, Steven J. Micheletti, Matthew H. McIntyre, Meghan E. Moreno, Priyanka Nandakumar, Dominique T. Nguyen, Jared O’Connell, Steve Pitts, G. David Poznik, Alexandra Reynoso, Shubham Saini, Morgan Schumacher, Leah Selcer, Anjali J. Shastri, Jingchunzi Shi, Suyash Shringarpure, Keaton Stagaman, Teague Sterling, Qiaojuan Jane Su, Joyce Y. Tung, Susana A. Tat, Vinh Tran, Xin Wang, Wei Wang, Catherine H. Weldon, Amy L. Williams, Peter Wilton.

